# Strategic infarct locations for post-stroke depressive symptoms: a lesion- and disconnection-symptom mapping study

**DOI:** 10.1101/2021.05.05.442398

**Authors:** Nick A. Weaver, Jae-Sung Lim, Janniek Schilderinck, Geert Jan Biessels, Yeonwook Kang, Beom Joon Kim, Hugo J. Kuijf, Byung-Chul Lee, Keon-Joo Lee, Kyung-Ho Yu, Hee-Joon Bae, J. Matthijs Biesbroek

**Affiliations:** Department of Neurology and Neurosurgery, UMC Utrecht Brain Center, Utrecht, the Netherlands; Department of Neurology, Asan Medical Center, Seoul, Republic of Korea; Department of Neurology, Hallym University Sacred Heart Hospital, Hallym Neurological Institute, College of Medicine, Hallym University, Anyang, Republic of Korea; Department of Psychology, Hallym University, Chuncheon, Republic of Korea; Department of Neurology, Seoul National University Bundang Hospital, Seoul National University College of Medicine, Seongnam, Republic of Korea; Image Sciences Institute, University Medical Center Utrecht, Utrecht, the Netherlands

**Keywords:** ischemic stroke, lesion-symptom mapping, lesion location, magnetic resonance imaging, post-stroke depression, Geriatric Depression Scale

## Abstract

**Background:** Depression is the most common neuropsychiatric complication after stroke. Infarct location is associated with post-stroke depressive symptoms (PSDS), but it remains debated which brain structures are critically involved. We performed a large-scale lesion-symptom mapping study to identify infarct locations, and white matter disconnections, associated with PSDS.

**Methods:** We included 553 patients (age 69±11 years, 42% female) with acute ischemic stroke. PSDS were measured using the 30-item Geriatric Depression Scale (GDS-30). Multivariable support vector regression (SVR)-based analyses were performed both at the level of individual voxels (SVR-VLSM) and predefined regions of interest (SVR-ROI) to relate infarct location to PSDS. We externally validated our findings in an independent stroke cohort (N=459). Finally, disconnectome-based analyses were performed using SVR-VLSM, in which white matter fibers disconnected by the infarct were analyzed instead of the infarct itself. **Results:** Infarcts in the right amygdala, right hippocampus and right pallidum were consistently associated with PSDS (permutation-based p<0.05) in SVR-VLSM and SVR-ROI. External validation (N=459) confirmed the association between infarcts in the right amygdala and pallidum, but not the right hippocampus, and PSDS. Disconnectome-based analyses revealed that disconnections in the right parahippocampal white matter, right thalamus and pallidum, and right anterior thalamic radiation were significantly associated (permutation-based p<0.05) with PSDS.

**Conclusions:** Infarcts in the right amygdala and pallidum, and disconnections of right limbic and frontal cortico-basal ganglia-thalamic circuits, are associated with PSDS. Our findings provide a comprehensive and integrative picture of strategic infarct locations for PSDS, and shed new light on pathophysiological mechanisms of depression after stroke.

## Introduction

Depression is the most common neuropsychiatric complication after stroke, affecting approximately one-third of stroke survivors (1). Individuals with post-stroke depression are at a higher risk of functional impairment, reduced quality of life, and increased mortality (2). Despite its high prevalence and debilitating impact on patients, the pathophysiology of post-stroke depression remains poorly understood (3). Better understanding of post-stroke depression pathophysiology is a critical step towards developing targeted prevention and treatment strategies, as was emphasized in a recent statement from the American Heart Association and American Stroke Association (3).

Infarct location has been identified as a contributor to post-stroke depressive symptoms (PSDS), yet it remains debated which brain structures are critically involved. Numerous reviews on this topic have been published in the past two decades, but results are inconsistent (see Supplementary Table 1). The most recent systematic review and meta-analysis to date (covering literature until 2016) found that infarcts in frontal and subcortical locations, and in the basal ganglia, were significantly associated with PSDS (4). However, many original studies only considered relatively crude spatial characteristics (e.g. hemispheric lateralization or frontal lobe involvement) (4,5). In the past decade, several studies have applied lesion-symptom mapping techniques to analyze the relationship between infarct location and depressive symptoms at a more detailed spatial resolution, usually at the level of individual brain voxels (6–10). Previous studies found significant associations for infarcts in the left dorsolateral prefrontal cortex (DLPFC) (7), left ventrolateral prefrontal cortex (11), and posterior cerebellum (8), while three studies found no significant associations (6,9,10). The lack of consistent evidence may be due to modest sample sizes in individual studies (i.e. largest single-center sample N=270), or specific inclusion criteria (e.g. only left hemispheric infarcts (7) or isolated cerebellar infarcts (8)), both resulting in limited lesion coverage of the brain. Lesion coverage is important in lesion-symptom mapping studies, because if an anatomical structure is not damaged in a sufficient number of patients, potential structure-function relations will not be detected. Of note, the largest lesion-symptom mapping study on PSDS to date (N=461 total) largely consisted of a mixed population, including individuals with traumatic brain injury and intracerebral hemorrhages and only ∼40% with ischemic stroke (9). In light of differences in disease mechanisms, this may have affected the results. A large-scale study with a homogeneous population could overcome these challenges.

Notably, a recent study proposed that disconnections caused by an infarct, rather than the location of the infarct itself, might also be related to depressive symptoms. In a combined sample from five datasets with different lesion etiologies, no specific lesion locations were associated with depressive symptoms, yet when functional disconnections (as identified using a normative functional connectome dataset) caused by each lesion were analyzed, robust associations with functional disconnection of the left DLPFC were found (9).

In this large-scale study we aimed to identify infarct locations associated with PSDS using multivariable lesion-symptom mapping, which we subsequently validated in an independent stroke cohort. Additionally, building upon emerging evidence, we performed structural disconnection-symptom mapping analyses using the same multivariable analysis approach.

## Methods and Materials

### Study population

Patients were selected from the Bundang Vascular Cognitive Impairment (VCI) cohort, consisting of patients admitted to Seoul National University Bundang Hospital, Republic of Korea, with acute ischemic stroke between 2007 and 2018 (12). A total of 553 patients were selected based on the following inclusion criteria: 1) brain magnetic resonance imaging (MRI) showing the acute symptomatic infarct(s) on diffusion-weighted imaging (DWI) and/or fluid-attenuated inversion recovery (FLAIR), 2) successful infarct segmentation and registration (section “Generation of lesion maps”), 3) no previous cortical infarcts, large subcortical infarcts or intracerebral hemorrhages on MRI (section “Generation of lesion maps”), and 4) available 30-item Geriatric Depression Scale (GDS-30) assessment, and clinical data on age, sex, and education. A flowchart of patient selection is provided in Figure 1.

**Figure 1.**
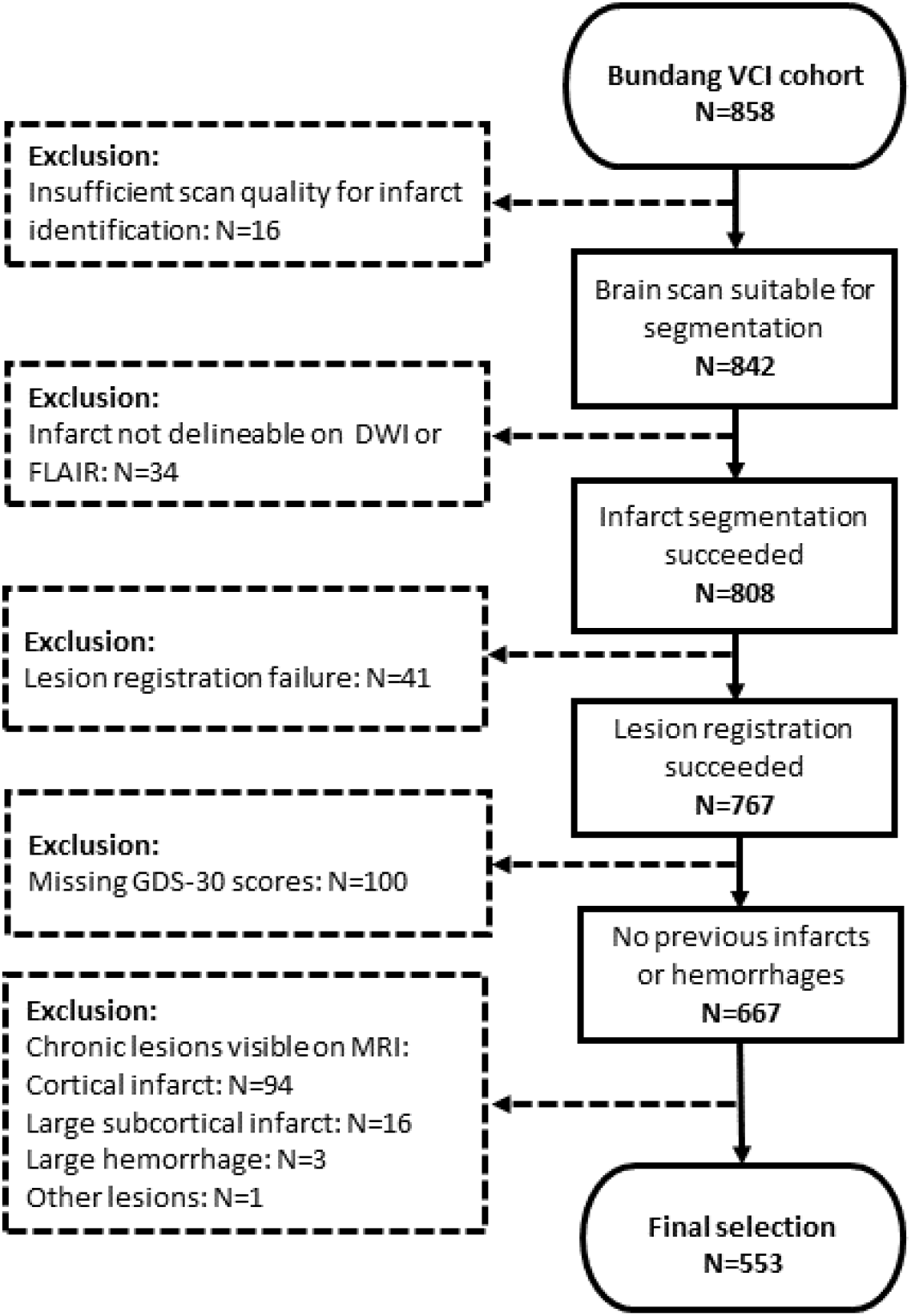
Flowchart of patient selection.

### Generation of lesion maps

Brain MRI, including DWI and FLAIR sequences, was performed with a 3 tesla MRI scanner in the first week after stroke onset. Details regarding the MRI protocol are provided in the Supplementary Material. Lesion data was available from a previous project (13). In short, infarct segmentation and subsequent registration to the T1 1 mm MNI-152 brain template (resolution 1×1×1 mm) (14) were performed in accordance with a previously published protocol (15). First, acute infarcts were manually segmented on DWI (N=536; 97%) or FLAIR (N=17; 3%) sequences using in-house developed software built in MeVisLab (MeVis Medical Solutions AG, Bremen, Germany) (16). ADC maps and T1 sequences were used as reference when available. Next, all scans and the corresponding lesion maps were transformed to MNI-152 space with the RegLSM tool (https://github.com/Meta-VCI-Map/RegLSM). Quality checks of all registration results were performed by an experienced rater (N.A.W.). Manual adaptations were made in case of minor registration errors (N=255; 46%). Presence of chronic cortical infarcts (any size), large subcortical infarcts (>15 mm), and large intracerebral hemorrhages (>10 mm) was assessed by an experienced rater (N.A.W.), and patients with any of these lesions were excluded from the current study.

### Generation of disconnectome maps

Disconnectome maps were calculated using the BCBtoolkit (17). This approach utilizes connectome maps derived from diffusion weighted imaging tractography data from 10 healthy controls (18) and uses the patient’s lesion maps as seed region for tractography to identify disconnected white matter fibers (17). In a previous validation study, these 10 healthy controls proved to be enough to obtain disconnectome maps that match the overall population (i.e. >70% shared variance) and to ensure that the reliability of these disconnection maps did not decrease with increasing age of the patient (17). For each healthy control, tractography was estimated as previously described (19). The lesion map in MNI-152 space of each patient was registered to the native space of the healthy controls using affine and diffeomorphic deformations (20,21) and subsequently used as seed for the tractography in Trackvis (22). Tractographies from the lesions were transformed into visitation maps (23), binarized and registered back to MNI-152 space using the inverse of precedent deformations. Finally, a percentage overlap map was generated for each patient by summing the normalized visitation map of each healthy subject for each voxel in MNI-152 space. Hence, in the resulting disconnectome map, the value in each voxel indicates the probability of disconnection from 0 to 100%. The disconnectome maps were subsequently dichotomized using a threshold of ≥50%. Thus, for each voxel in the brain, the binary disconnectome maps indicated whether an anatomical connection with the lesioned area would normally exist in healthy individuals, and consequently whether a voxel would become disconnected by the lesion.

### Assessment of post-stroke depressive symptoms

Depressive symptoms were measured using a validated Korean version of the GDS-30 (24). The GDS is a self-report questionnaire on depressive symptoms; patients respond to a list of questions in “yes/no” format, with higher scores indicating a larger number of depressive symptoms (25). The GDS-30 was administered within the first year post-stroke by trained clinical neuropsychologists who were blinded to patients’ clinical and neuroradiological profiles. At the discretion of the attending physician, patients were excluded if they suffered from disabilities that would interfere with neuropsychiatric testing, including neurological deficits such as severe aphasia or severe motor weakness, or impairment of hearing or vision.

### Statistical analyses

#### Outcome measures

GDS-30 scores were converted into standardized z-scores and multiplied by -1, so that lower scores indicated more depressive symptoms. Correction for covariates was performed in two steps. First, z-scores were corrected for age, sex, and years of education using linear regression. Second, we further took the influence of stroke severity, physical disability, and cognitive impairment into account, because these were identified as most consistent predictors of PSDS in a recent review (3). For this purpose, we performed additional correction for National Institutes of Health Stroke Scale (NIHSS) score, impairment of Activities of Daily Living (ADL) according to the Korean Instrumental Activities of Daily Living, and presence of post-stroke cognitive impairment (PSCI) (definitions provided in Supplementary Material). Both corrected z-scores (i.e. corrected for 3 and 6 factors respectively) were analyzed separately as measures for PSDS in all SVR-based analyses mentioned below.

#### Lesion-symptom mapping

We performed multivariable support vector regression-based (SVR) analyses to determine the association between infarct location and post-stroke depressive symptoms. Two independent, hypothesis-free approaches were applied in accordance with previously published methods (26,27): voxel-based (SVR-VLSM) and region of interest-based lesion-symptom mapping (SVR-ROI). Both SVR-based methods were applied because of their complementary strengths: SVR-VLSM offers a much higher spatial resolution, while SVR-ROI takes the cumulative lesion burden within predefined brain regions into account. An important advantage of these multivariable methods is that the interrelation between voxels and ROIs is taken into account, thereby providing a higher spatial accuracy than mass-univariable methods (27).

SVR-VLSM was performed using Python (SciPy 1.4.1). Only voxels damaged in at least 5 patients were included in the analyses. A linear SVR model with feature selection was used, in accordance with previous studies (26,27). In the feature selection step, only voxels in which the presence of a lesion was univariately associated with depressive symptoms (two-sided t-test, uncorrected p<0.05) were selected to reduce noise. Next, parameter training of the linear SVR model was performed to determine the optimal regularization parameter (C) and epsilon to maximize the prediction performance of the z-scores. The prediction performance was calculated for each combination of C and epsilon values by determining the mean Pearson correlation coefficient between the real and predicted z-scores with 5-fold cross-validations (optimal parameters in Supplementary Table 2). Statistical inference was performed by shuffling the observations of z-scores to create pseudo weight coefficients, and the significance level of each voxel was calculated by counting the number of pseudo weights larger than the real weight in 5000 permutations. Voxels with permutation-based p<0.05 were treated as statistically significant. We corrected for total infarct volume by weighting the lesioned voxels in inverse proportion to the square root of total infarct volume prior to model training.

SVR-ROI was performed using MATLAB (version R2018a). ROIs were defined using the AAL atlas (119 ROIs) (28) and ICBM-DTI-81 white matter tract atlas (50 ROIs) (29,30) in MNI-152 space (14). Infarct volumes for each ROI were calculated in milliliters. Only ROIs damaged in at least 5 patients were included in the analyses. Similar to the SVR-VLSM, a linear SVR model with feature selection was used (31). In the feature selection step, ROIs with a univariate significant Pearson correlation (p<0.05) between infarct volume and PSDS were selected. The parameter training and statistical inference of SVR-ROI corresponded with the SVR-VLSM (optimal parameters in Supplementary Table 3). ROIs with permutation-based p<0.05 were treated as statistically significant. We corrected for total infarct volume by including it as covariate alongside the ROI volumes in the SVR models.

#### Disconnection-symptom mapping

The association between anatomical disconnections and depressive symptoms was analyzed at the level of individual voxels using SVR-VLSM. The same SVR-VLSM approach was applied (see section “Lesion-symptom mapping”), but with two differences: disconnectome maps were entered as independent variable instead of lesion maps, and we corrected for total disconnectome volume instead of total infarct volume. Voxels with permutation-based p<0.05 were treated as statistically significant.

### External validation

To determine the external validity and reproducibility of our findings, we determined whether the statistically significant ROIs identified in the SVR-ROI analysis were also associated with depressive symptoms in an independent study sample from the Hallym VCI cohort (N=459) (32). Study protocols and patient characteristics were comparable to the Bundang VCI cohort, and lesion data was collectively prepared in a previous collaboration between the two centers (13). In the Hallym VCI cohort, depressive symptoms were measured using a validated Korean version of the 15-item Geriatric Depression Scale (GDS-15) (33), which is a shorter version of the GDS-30 questionnaire. Details on patient characteristics and selection for the validation sample are provided in the Supplementary Material. We followed the same data processing steps as for the Bundang VCI cohort: GDS-15 scores were converted into standardized z-scores and multiplied by -1, and infarct volumes per ROI were calculated using the AAL atlas (28) and ICBM-DTI-81 atlases (29,30) in MNI-152 space (14). Significant ROIs were selected from the SVR-ROI results (p<0.05), and these ROI volumes were entered in separate multiple linear regression model together with age, sex, and education. This was done for both Bundang VCI and Hallym VCI datasets to allow for direct comparison. As negative control, models were also built for three ROIs that were selected based on the following criteria: 1) no association with PSDS in any of the analyses using GDS-30 in the main sample (NB: if an ROI was significantly associated with PSDS in one hemisphere, the contralateral ROI was also excluded); 2) 1 left supratentorial, 1 right supratentorial, and 1 infratentorial ROI were randomly selected to ascertain spatial independence of the selected negative control ROIs. All ROI volumes underwent cube root transformation to meet the normality assumptions of multiple linear models. Standardized betas (Stβ) were calculated for each ROI.

### Ethics statement

Patient data was collected in accordance with study protocols approved by the Institutional Review Boards of the Seoul National University Bundang Hospital and Hallym University Sacred Heart Hospital with a waiver of patient consent because of the retrospective nature of the study and minimal risk to the study participants. Use of the acquired data for the current study was also approved by these Institutional Review Boards.

## Results

### Study population

Demographic and clinical characteristics of the 553 included patients are provided in Table 1. Mean age was 69 years (SD=11), 42% were female (N=233/553), and median years of education was 9 (IQR=6-14). GDS-30 scores ranged from 0 to 30 (i.e. full range of scores), with a median of 13 (IQR=7-20). PSCI and ADL impairment occurred in 58% and 25% of patients respectively, and median NIHSS was 3 (IQR=1-5). GDS-30 and cognitive assessments generally took place in the first 6 months post-stroke, with a median time interval of 105 days (IQR=11-168). Infarcts were generally small, with a median normalized volume of 3.3 mL (IQR=1.1-15.1).

**Table 1.** Demographic and clinical characteristics of study sample (N=553)

| **Demographics and clinical characteristics** | **Total sample (N=553)** |
| --- | --- |
| Age (years), mean (SD) | 69.0 (11.0) |
| Female, N (%) | 233 (42%) |
| Years of education, median (IQR) | 9 (6-14) |
| NIHSS at admission, median (IQR) | 3 (1-5) |
| K-IADL score, median (IQR) | 0.10 (0.00-0.42),<br>(missing N=4) |
| ADL impairment (K-IADL score > 0.43) | 137 (25%)<br>(missing N=4) |
| **Hand preference, N (%)** | (missing N=2) |
| Right | 534 (97%) |
| Left | 6 (1%) |
| Ambidextrous | 11 (2%) |
| **Vascular risk factors, N (%)** |  |
| Hypertension | 422 (76%) |
| Hyperlipidemia | 135 (24%) |
| Current smoker | 119 (22%) |
| Past smoker | 113 (20%) |
| Diabetes Mellitus | 179 (32%) |
| Atrial fibrillation | 84 (15%)<br>(missing N=27) |
| **Neuropsychiatric and cognitive assessment** |  |
| Time interval between stroke onset and neuropsychiatric and cognitive assessment (days), median (IQR), range | 105 (11-168),<br>range 1-361 |
| GDS-30 scores, median (IQR), range | 13 (7-20), range 0-30 |
| Presence of post-stroke cognitive impairment, N (%) | 323 (58%)<br>(missing N=10) |
| **Brain MRI** |  |
| Time interval between stroke onset and MRI (days), median (IQR), range | 5 (4-6), range 0-41 |
| Total infarct volume in ml*, median (IQR), range | 3.3 (1.1-15.1),<br>range 0.04-535.1 |

Missing data is noted behind the respective variable, when appropriate. Valid percent is indicated in cases with missing data.
\*Normalized volumes after registration to the MNI-152 template. Abbreviations: IQR, interquartile range; MRI, magnetic resonance imaging; SD, standard deviation.

### Lesion-symptom mapping results

Infarct distribution was symmetrical and subcortical regions were more commonly damaged than cortical regions (Figure 2A). Because of the large sample size, brain coverage was relatively high. In the SVR-VLSM analyses, 52% of voxels of the MNI-152 template (944,338/1,827,240 voxels) were included. Parts of the midbrain, temporal lobes, cerebellum, and anterior cerebral artery territory could not be included due to infrequent involvement (i.e. damaged in <5 patients). In the SVR-ROI analyses, 167 out of 169 ROIs were included.

**Figure 2.**
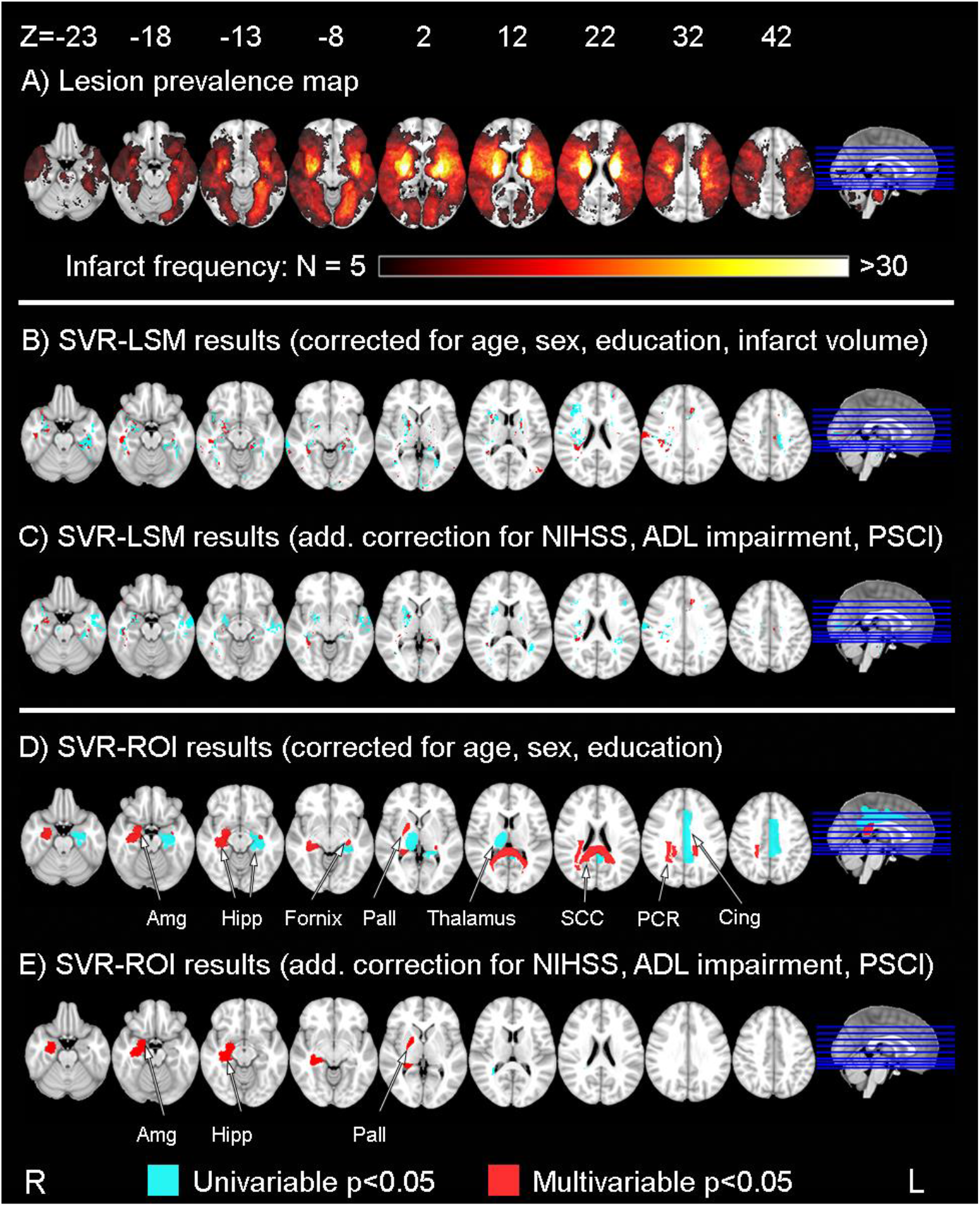
Lesion prevalence map and lesion-symptom mapping results (N=553) Results are depicted on the Montreal Neurological Institute 152 T1 1mm template (Fonov et al. 2011). (A) Infarct prevalence map showing voxels that are damaged in ≥5 patients (total N=553). Only colored voxels were included in subsequent analyses. (B-C) Results of voxel-based lesion-symptom mapping (SVR-VLSM). First, feature selection was performed using univariable voxel-based lesion-symptom mapping (two-sample t-test; p<0.05). Next, linear SVR-VLSM was performed on these selected voxels (shown in red and cyan). Significant voxels from the SVR-LSM analysis are shown in red (p<0.05 based on 5000 permutations); feature-selected but non-significant voxels are shown in cyan. (D-E) Results of region of interest-based lesion-symptom mapping (SVR-ROI). ROIs were defined by the AAL grey matter atlas and ICBM-DTI-81 white matter tract atlas in MNI-152 space. First, feature selection was performed using regional infarct volumes and total infarct volume (Pearson correlation; p<0.05). Note that total infarct volume was not selected in this step. Next, linear SVR-ROI was performed on these selected ROIs (shown in red and cyan). Significant ROIs from the SVR-ROI analysis are shown in red (p<0.05 based on 5000 permutations); feature-selected but non-significant voxels are shown in cyan. Abbreviations: NIHSS = National Institutes of Health Stroke Scale, ADL = Activities of Daily Living, PSCI = post-stroke cognitive impairment, Amg = amygdala, Hipp = hippocampus, Pall = pallidum, SCC = splenium of the corpus callosum, PCR = posterior corona radiata, Cing = cingulate gyrus.

SVR-VLSM results are shown in Figure 2B-C. Voxel clusters in the right amygdala, right pallidum, right corona radiata, right sagittal striatum, bilateral hippocampi, bilateral fornices, and left cingulum of the hippocampus were significantly associated with more PSDS (permutation-based p<0.05), after correction for age, sex, education, total infarct volume, NIHSS score, ADL impairment and PSCI (Figure 2C and Supplementary Table 4). Voxel clusters in the left pallidum and right postcentral gyrus were also associated with more PSDS after correction for age, sex, education and total infarct volume (Figure 2B), but no longer statistically significant after additional correction for NIHSS score, ADL impairment and PSCI (Figure 2C). SVR-ROI results are shown in Figure 2D-E. Total infarct volume was not univariately associated with more PSDS, therefore it was not included in the SVR-ROI models as covariate. Regional infarct volumes in the right amygdala, right pallidum and right hippocampus were significantly associated with more PSDS (permutation-based p<0.05), after correction for age, sex, education, NIHSS score, ADL impairment and PSCI (Figure 2E). Regional infarct volumes in the left fornix, splenium of the corpus callosum, and right posterior corona radiata were also associated with more PSDS after correction for age, sex, education (Figure 2D), but no longer statistically significant after additional correction for NIHSS score, ADL impairment and PSCI (Figure 2E). No infarct locations were significantly associated with less PSDS in either SVR-VLSM or SVR-ROI analyses.

### Disconnection-symptom mapping results

Disconnections of nearly all supratentorial white matter fibers were included in the analyses, with a symmetrical distribution (Figure 3A). Disconnectome-based analyses with SVR-VLSM revealed that disconnections in the right parahippocampal white matter, right thalamus and pallidum, and right anterior thalamic radiation were significantly associated (p<0.05) with more PSDS, after correction for age, sex, education, total disconnectome volume, NIHSS score, ADL impairment and PSCI (Figure 3B).

**Figure 3.**
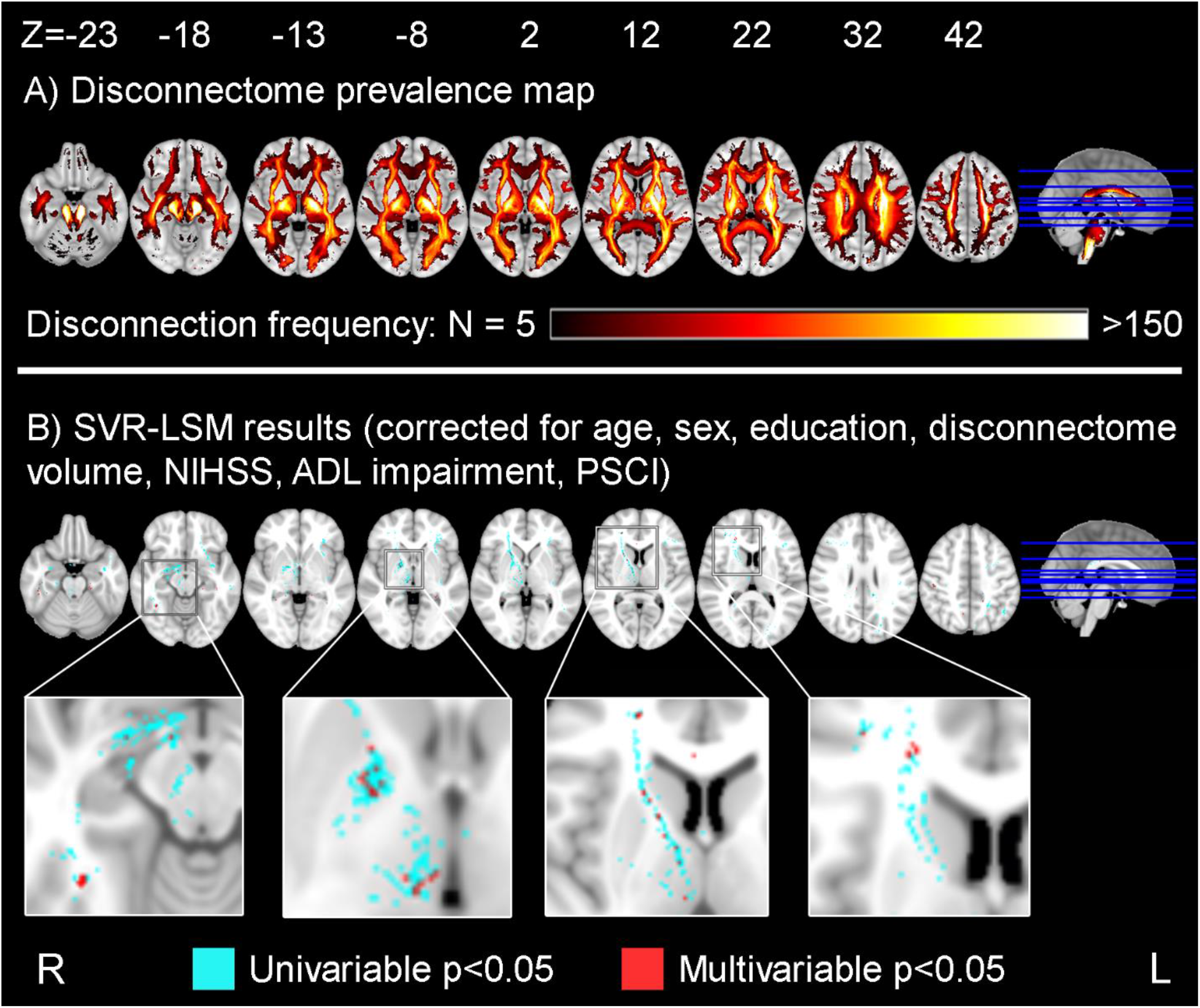
Structural disconnection prevalence map and disconnection-symptom mapping results. Results are depicted on the Montreal Neurological Institute 152 T1 1mm template (Fonov et al. 2011). (A) Structural disconnection prevalence map showing voxels that are affected in ≥5 patients (total N=553). Only colored voxels were included in subsequent analyses. (B) Results of voxel-based disconnection-symptom mapping (SVR-LSM). Note that this method is the same as for the infarct location-based analysis (see Figure 2B-C), but using the map of disconnections caused by the infarct as determinant instead of the infarct itself. First, feature selection was performed using univariable voxel-based lesion-symptom mapping (two-sample t-test; p<0.05). Next, linear SVR-LSM was performed on these selected voxels (shown in red and cyan). Significant voxels from the SVR-LSM analysis are shown in red (p<0.05 based on 5000 permutations); feature-selected but non-significant voxels are shown in cyan. Abbreviations: NIHSS = National Institutes of Health Stroke Scale, ADL = Activities of Daily Living, PSCI = post-stroke cognitive impairment.

### External validation

Demographic and clinical characteristics of the validation sample (Hallym VCI, N=459) are shown in Supplementary Table 5. Compared to the main sample (Bundang VCI, N=553), the validation sample was younger (65 versus 69 years), had smaller infarcts (median infarct volume: 1.9 versus 3.3 mL), and had a lower occurrence of PSCI (44% versus 58%) and ADL impairment (13% versus 25%). Brain lesion coverage was lower in the validation sample, particularly in the bilateral temporal and occipital lobes, due to the smaller infarcts and smaller sample size (Supplementary Figure 2). Based on the SVR-ROI results (Figure 2E), the right amygdala, right hippocampus, and right pallidum were selected as ROIs for the external validation (Table 2). In the validation sample, regional infarct volumes in the right amygdala (Stβ=- 0.15, p=0.001) and right pallidum (Stβ=-0.14, p=0.002) were significantly associated with more PSDS, with slightly larger effect sizes than in the main sample (Stβ=-0.11 and -0.09 respectively). By contrast, the association with the right hippocampus was not confirmed (Stβ=-0.03, p=0.5). ROIs randomly selected as negative controls were not associated with PSDS in the validation sample (Table 2).

**Table 2.** External validation of identified regions of interest in an independent dataset.

| **Model: age , sex, education, and ROI volume** | **Bundang VCI (N=553)** |  |  | **Hallym VCI (N=459)** |  |  |
| --- | --- | --- | --- | --- | --- | --- |
|  | **Original sample** |  |  | **Validation sample** |  |  |
|  | **Standardized B for ROI volume** | **p-value** | **N patients with ROI damaged** | **Standardized B for ROI volume** | **p-value** | **N patients with ROI damaged** |
| Right amygdala | -0.11 | 0.007 | 37 | -0.15 | 0.001 | 32 |
| Right hippocampus | -0.08 | 0.043 | 57 | -0.03 | 0.478 | 38 |
| Right pallidum | -0.09 | 0.023 | 65 | -0.14 | 0.002 | 67 |
| Left insula (negative control)* | 0.01 | 0.800 | 128 | -0.03 | 0.478 | 85 |
| Middle cerebellar peduncle (negative control)* | 0.01 | 0.847 | 101 | 0.01 | 0.821 | 77 |
| Right middle occipital gyrus (negative control)* | -0.03 | 0.480 | 56 | -0.07 | 0.113 | 39 |

\*Models were also built for three ROIs that were not associated with PSDS in the Bundang VCI cohort to strengthen the generalizability of the findings.

## Discussion

In this large-scale lesion-symptom mapping study, we newly identified the right amygdala as infarct location associated with PSDS, and further pinpointed the right pallidum as key location within the basal ganglia. This finding was replicated in an independent cohort. Furthermore, we showed that structural disconnections located in right thalamo-cortical and parahippocampal white matter were associated with PSDS. This converging evidence demonstrates that infarcts and resulting structural disconnections located in right frontal cortico-thalamic and limbic circuits are related to PSDS.

The relationship between infarct location and PSDS has been extensively studied, yet previous findings varied widely. Early studies were limited by technical constraints and could only consider relatively crude spatial characteristics, such as hemispheric lateralization or lobar involvement (5). Although recent lesion-symptom mapping studies provided a higher spatial resolution, they still lacked consistent evidence, likely due to modest sample sizes and subsequent limited lesion coverage and inclusion of mixed pathologies. In our large homogeneous sample we achieved high brain coverage (i.e. 52% compared to <25% in most monocenter lesion-symptom mapping studies (26)) and performed state-of-the-art analyses at a detailed spatial resolution, and thereby created the most comprehensive picture of strategic infarct locations for PSDS to date.

A recent systematic review and meta-analysis found that infarcts in the basal ganglia were significantly associated with PSDS (4). We confirm and extend these findings by pinpointing the pallidum, and more specifically the ventral pallidum (i.e. based on the VLSM analyses), as key anatomical structure within the basal ganglia. This increased anatomical detail is important, because the basal ganglia consist of a multitude of structures and projections, within which the ventral pallidum is a core component of the limbic loop (34). Furthermore, we identified the right amygdala as strategic infarct location, which has only been suggested once before in a study in 68 stroke patients (35). This limited prior evidence on the amygdala might be due to its infrequent involvement and subsequent limited statistical power in smaller studies, considering that the amygdala is supplied by the anterior choroidal artery (36) which only accounts for 8% of patients with acute ischemic stroke (37). An important strength of our study is that we externally validated the relevance of these locations in an independent dataset. Despite the use of a different version of the GDS and differences in cohort characteristics (i.e. age, infarct size, and presence of functional impairment), the right amygdala and pallidum showed comparable associations with PSDS in both stroke cohorts. This demonstrates the robustness and generalizability of our findings.

Recent evidence suggests that PSDS can result from interruptions of functional brain networks caused by infarcts (9). Our complementary lesion- and disconnection-symptom mapping approaches provide an integrated perspective, showing that both infarct location and structural disconnections are important. Our voxel-based analyses showed a clear pattern of involvement of voxel clusters in limbic system structures, including the amygdala and ventral pallidum, but also the hippocampus, fornix, cingulate gyrus, and cingulum of the hippocampus. Meanwhile, the disconnetome-based analysis showed specific involvement of tracts connecting the right frontal cortex to the pallidum and thalamus, and from the thalamus to the parahippocampal white matter. Taking these findings together, the right limbic and frontal cortico-basal ganglia-thalamic circuits consistently emerge as key brain networks, in which both direct damage to key brain structures and disruption of connecting pathways are linked to PSDS.

The pathophysiology of PSDS is complex and involves a combination of biological and psychosocial factors (3). Our findings strengthen the notion that lesion location is a contributing biological factor, and suggest a central role of the limbic system. This is in line with evidence from studies on depression as primary psychiatric disorder. Decreased amygdala and hippocampus volumes were found in patients with depression, and functional MRI and PET studies have reported changes in metabolism or activity in the amygdala and hippocampus (38). Diffusion tensor imaging-based studies have reported structural alterations to fronto-striato-thalamic white matter tracts in patients with depression (39). Meanwhile, white matter hyperintensities are also thought to contribute to depressive symptoms in elderly people through disruption of structural brain networks (i.e. following the “vascular depression” hypothesis) (40,41). White matter hyperintensities located in (pre)frontal and temporal regions have been associated with depressive symptoms (42–45), which aligns with locations identified in our disconnectome-based analyses. This suggests that vascular damage to the fronto-limbic circuits represents a common mechanism of depressive symptoms in cerebral small vessel disease and acute ischemic stroke (5).

Post-stroke physical and cognitive deficits have also been linked to PSDS, indirectly suggesting that PSDS may be a psychological reaction to these deficits (3,46). Our results were independent of measures of stroke severity, physical disability and cognitive impairment, which supports the notion of a direct effect of infarcts on the depressive symptoms.

A notable observation is that most of the infarct locations associated with PSDS were located in the right hemisphere, even though our lesion coverage of the right and left hemisphere was nearly symmetrical. The impact of lesion laterality in PSDS has been a topic of extensive discussion in the field, but no clear pattern has been found. A recent meta-analysis found no association with laterality in 60 studies, neither in the pooled analysis nor in subgroup analyses stratified by study phase (i.e. acute, subacute, and chronic) (4). Evidence on lateraterality from other neuroimaging research on depression (e.g. EEG, fMRI, PET) is also inconclusive (47). Our results should therefore not be interpreted as specific to the right hemisphere, but rather to highlight the importance of specific brain structures and networks.

Some limitations to our study must also be noted. First, the GDS-30 was designed as a screening tool. While the questionairre allows us to reliably quantify the number of depressive symptoms, it is not suitable for determining a formal diagnosis of post-stroke depression. A formal diagnosis might be more relevant from a clinical perspective, but would require elaborate evaluation by a psychiatrist, which might prove difficult to achieve in large-scale stroke studies. Second, information on clinical history of depression was not collected, therefore we could not take potential influence of pre-stroke symptoms into account. Third, our analysis of structural disconnections was based on a brain connectivity template, and not the actual brain connectivity of individual patients, therefore patient-specific variability in brain connectivity might not be fully accounted for.

Our study contributes to the knowledge of the pathophysiology of post-stroke depression by highlighting the involvement of specific brain structures and connections (3). As a next step, the value of strategic infarct location as a prognostic imaging marker could be assessed using predictive modeling, similar to a recent study on PSCI prediction (48). If successful, this could facilitate early identification of patients at risk of developing PSDS, permit more timely intervention and treatment of these debilitating symptoms, and thereby lower the burden on both patients and caregivers.

## Acknowledgements

This work was supported by Vici grant 918.16.616 from ZonMw, The Netherlands, Organization for Health Research. J.M.B.’s work was supported by a Young Talent Fellowship from the UMC Utrecht Brain Center, The Netherlands. Data from this article was previously submitted as an abstract for the European Stroke Organization conference 2021. The authors thank Gözdem Arikan and Angelina Kancheva for their help with the infarct segmentations, and Jaeseol Park and Eunbin Ko for their help in organizing the clinical data.

## Disclosures

Nick A. Weaver, Jae-Sung Lim, Janniek Schilderinck, Geert Jan Biessels, Yeonwook Kang, Beom Joon Kim, Hugo J. Kuijf, Byung-Chul Lee, Keon-Joo Lee, Kyung-Ho Yu, Hee-Joon Bae, and J. Matthijs Biesbroek reported no biomedical financial interests or potential conflicts of interest.

## Notes

### Competing Interest Statement

The authors have declared no competing interest.

## References

1. Hackett ML, Pickles K (2014): Part I: Frequency of depression after stroke: An updated systematic review and meta-analysis of observational studies. Int J Stroke 9: 1017–1025.

2. Kutlubaev MA, Hackett ML (2014): Part II: Predictors of depression after stroke and impact of depression on stroke outcome: An updated systematic review of observational studies. Int J Stroke 9: 1026–1036.

3. Towfighi A, Ovbiagele B, El Husseini N, Hackett ML, Jorge RE, Kissela BM, et al. (2017, February 1): Poststroke Depression: A Scientific Statement for Healthcare Professionals from the American Heart Association/American Stroke Association. Stroke, vol. 48. Lippincott Williams and Wilkins, pp e30–e43.

4. Douven E, Köhler S, Rodriguez MMF, Staals J, Verhey FRJ, Aalten P (2017, September 1): Imaging Markers of Post-Stroke Depression and Apathy: a Systematic Review and Meta-Analysis. Neuropsychology Review, vol. 27. Springer New York LLC, pp 202–219.

5. Nickel A, Thomalla G (2017, September 21): Post-stroke depression: Impact of lesion location and methodological limitations-a topical review. Frontiers in Neurology, vol. 8. Frontiers Media S.A. https://doi.org/10.3389/fneur.2017.00498

6. Gozzi SA, Wood AG, Chen J, Vaddadi K, Phan TG (2014): Imaging predictors of poststroke depression: Methodological factors in voxel-based analysis. BMJ Open 4: 4948.

7. Grajny K, Pyata H, Spiegel K, Lacey EH, Xing S, Brophy C, Turkeltaub PE (2016): Depression symptoms in chronic left hemisphere stroke are related to dorsolateral prefrontal cortex damage. J Neuropsychiatry Clin Neurosci 28: 292–298.

8. Kim NY, Lee SC, Shin JC, Park JE, Kim YW (2017): Voxel-based lesion symptom mapping analysis of depressive mood in patients with isolated cerebellar stroke: A pilot study. NeuroImage Clin 13: 39–45.

9. Padmanabhan JL, Cooke D, Joutsa J, Siddiqi SH, Ferguson M, Darby RR, et al. (2019): A Human Depression Circuit Derived From Focal Brain Lesions. Biol Psychiatry 86: 749–758.

10. Sagnier S, Munsch F, Bigourdan A, Debruxelles S, Poli M, Renou P, et al. (2019): The Influence of Stroke Location on Cognitive and Mood Impairment. A Voxel-Based Lesion-Symptom Mapping Study. J Stroke Cerebrovasc Dis 28: 1236–1242.

11. Klingbeil J, Brandt M-L, Wawrzyniak M, Stockert A, Schneider HR, Baum P, et al. (2021): Association of Lesion Location and Depressive Symptoms Poststroke. Stroke 52: 830–837.

12. Yu KH, Cho SJ, Oh MS, Jung S, Lee JH, Shin JH, et al. (2013): Cognitive impairment evaluated with vascular cognitive impairment harmonization standards in a multicenter prospective stroke cohort in Korea. Stroke 44: 786–788.

13. Weaver NA, Kancheva AK, Lim J-S, Biesbroek JM, Wajer IMH, Kang Y, et al. (2021): Post-stroke cognitive impairment on the Mini-Mental State Examination primarily relates to left middle cerebral artery infarcts. Int J Stroke 174749302098455.

14. Fonov V, Evans AC, Botteron K, Almli CR, McKinstry RC, Collins DL, Brain Development Cooperative Group (2011): Unbiased average age-appropriate atlases for pediatric studies. Neuroimage 54: 313–327.

15. Biesbroek JM, Kuijf HJ, Weaver NA, Zhao L, Duering M, Biessels GJ (2019): Brain Infarct Segmentation and Registration on MRI or CT for Lesion-symptom Mapping. J Vis Exp. https://doi.org/10.3791/59653

16. Ritter F, Boskamp T, Homeyer A, Laue H, Schwier M, Link F, Peitgen H (2011): Medical Image Analysis. IEEE Pulse 2: 60–70.

17. Foulon C, Cerliani L, Kinkingnéhun S, Levy R, Rosso C, Urbanski M, et al. (2018): Advanced lesion symptom mapping analyses and implementation as BCBtoolkit. GigaScience. https://doi.org/10.1093/gigascience/giy004

18. Rojkova K, Volle E, Urbanski M, Humbert F, Dell’Acqua F, Thiebaut de Schotten M (2016): Atlasing the frontal lobe connections and their variability due to age and education: a spherical deconvolution tractography study. Brain Struct Funct. https://doi.org/10.1007/s00429-015-1001-3

19. De Schotten MT, Dell’Acqua F, Forkel SJ, Simmons A, Vergani F, Murphy DGM, Catani M (2011): A lateralized brain network for visuospatial attention. Nat Neurosci. https://doi.org/10.1038/nn.2905

20. Klein A, Ghosh S, Avants B, Yeo B, Neuroimage BF-, 2010 undefined (n.d.): Evaluation of volume-based and surface-based brain image registration methods. Elsevier.

21. Avants BB, Tustison NJ, Song G, Cook PA, Klein A, Gee JC (2011): A reproducible evaluation of ANTs similarity metric performance in brain image registration. Neuroimage. https://doi.org/10.1016/j.neuroimage.2010.09.025

22. Wang R, Benner T (2007): Diffusion toolkit: a software package for diffusion imaging data processing and tractography. Proc Intl Soc Mag Reson Med.

23. Thiebaut de Schotten M, ffytche DH, Bizzi A, Dell’Acqua F, Allin M, Walshe M, et al. (2011): Atlasing location, asymmetry and inter-subject variability of white matter tracts in the human brain with MR diffusion tractography. Neuroimage. https://doi.org/10.1016/j.neuroimage.2010.07.055

24. Jung IK, Kwak D Il, Shin DK, Lee MS, Lee HS, Kim JY (1997): A Reliability and Validity Study of Geriatric Depression Scale. J Korean Neuropsychiatr Assoc 36: 103–112.

25. Yesavage JA, Brink TL, Rose TL, Lum O, Huang V, Adey M, Leirer VO (1982): Development and validation of a geriatric depression screening scale: A preliminary report. J Psychiatr Res 17: 37–49.

26. Weaver NA, Zhao L, Biesbroek JM, Kuijf HJ, Aben HP, Bae H-J, et al. (2019): The Meta VCI Map consortium for meta-analyses on strategic lesion locations for vascular cognitive impairment using lesion-symptom mapping: Design and multicenter pilot study. Alzheimer’s Dement Diagnosis, Assess Dis Monit 11: 310–326.

27. Zhao L, Biesbroek JM, Shi L, Liu W, Kuijf HJ, Chu WWC, et al. (2017): Strategic infarct location for post-stroke cognitive impairment: A multivariate lesion-symptom mapping study. J Cereb Blood Flow Metab. https://doi.org/10.1177/0271678X17728162

28. Tzourio-Mazoyer N, Landeau B, Papathanassiou D, Crivello F, Etard O, Delcroix N, et al. (2002): Automated Anatomical Labeling of Activations in SPM Using a Macroscopic Anatomical Parcellation of the MNI MRI Single-Subject Brain. Neuroimage 15: 273–289.

29. Mori S, Oishi K, Jiang H, Jiang L, Li X, Akhter K, et al. (2008): Stereotaxic white matter atlas based on diffusion tensor imaging in an ICBM template. Neuroimage. https://doi.org/10.1016/j.neuroimage.2007.12.035

30. Oishi K, Zilles K, Amunts K, Faria A, Jiang H, Li X, et al. (2008): Human brain white matter atlas: Identification and assignment of common anatomical structures in superficial white matter. Neuroimage. https://doi.org/10.1016/j.neuroimage.2008.07.009

31. Yourganov G, Fridriksson J, Rorden C, Gleichgerrcht E, Bonilha L (2016): Multivariate Connectome-Based Symptom Mapping in Post-Stroke Patients: Networks Supporting Language and Speech. J Neurosci 36: 6668– 6679.

32. Lim J, Kim N, Jang MU, Han M, Kim S, Baek MJ, et al. (2014): Cortical Hubs and Subcortical Cholinergic Pathways as Neural Substrates of Poststroke Dementia. https://doi.org/10.1161/STROKEAHA.

33. Cho MJ, Bae JN, Suh GH, Hahm BJ, Kim JK, Lee DW, Kang MH (1999): Validation of Geriatric Depression Scale, Korean Version(GDS) in the Assessment of DSM-III-R Major Depression. J Korean Neuropsychiatr Assoc 38: 48–63.

34. Smith KS, Tindell AJ, Aldridge JW, Berridge KC (2009, January 23): Ventral pallidum roles in reward and motivation. Behavioural Brain Research, vol. 196. Elsevier, pp 155–167.

35. Terroni L, Amaro E, Iosifescu D V., Tinone G, Sato JR, Leite CC, et al. (2011): Stroke lesion in cortical neural circuits and post-stroke incidence of major depressive episode: A 4-month prospective study. World J Biol Psychiatry 12: 539–548.

36. Kiernan JA (2012): Anatomy of the Temporal Lobe. Epilepsy Res Treat 2012: 1–12.

37. Ois A, Cuadrado-Godia E, Solano A, Perich-Alsina X, Roquer J (2009): Acute ischemic stroke in anterior choroidal artery territory. J Neurol Sci 281: 80–84.

38. Pandya M, Altinay M, Malone DA, Anand A (2012, December): Where in the brain is depression? Current Psychiatry Reports, vol. 14. NIH Public Access, pp 634–642.

39. Chen G, Hu X, Li L, Huang X, Lui S, Kuang W, et al. (2016): Disorganization of white matter architecture in major depressive disorder: A meta-analysis of diffusion tensor imaging with tract-based spatial statistics. Sci Rep 6. https://doi.org/10.1038/srep21825

40. Alexopoulos GS, Meyers BS, Young RC, Campbell S, Silbersweig D, Charlson M (1997): “Vascular depression” hypothesis. Archives of General Psychiatry, vol. 54. American Medical Association, pp 915–922.

41. Sneed JR, Culang-Reinlieb ME (2011): The vascular depression hypothesis: An update. American Journal of Geriatric Psychiatry, vol. 19. Elsevier B.V., pp 99–103.

42. Leeuwis AE, Weaver NA, Biesbroek JM, Exalto LG, Kuijf HJ, Hooghiemstra AM, et al. (2019): Impact of white matter hyperintensity location on depressive symptoms in memory-clinic patients: A lesion–symptom mapping study. J Psychiatry Neurosci 44. https://doi.org/10.1503/jpn.180136

43. Soennesyn H, Oppedal K, Greve OJ, Fritze F, Auestad BH, Nore SP, et al. (2012): White Matter Hyperintensities and the Course of Depressive Symptoms in Elderly People with Mild Dementia. Dement Geriatr Cogn Dis Extra 2: 97–111.

44. Dalby RB, Frandsen J, Chakravarty MM, Ahdidan J, Sørensen L, Rosenberg R, et al. (2010): Depression severity is correlated to the integrity of white matter fiber tracts in late-onset major depression. Psychiatry Res Neuroimaging 184: 38–48.

45. Sheline YI, Price JL, Vaishnavi SN, Mintun MA, Barch DM, Epstein AA, et al. (2008): Regional white matter hyperintensity burden in automated segmentation distinguishes late-life depressed subjects from comparison subjects matched for vascular risk factors. Am J Psychiatry 165: 524–532.

46. Nys GMS, Van Zandvoort MJE, Van Der Worp HB, De Haan EHF, De Kort PLM, Kappelle LJ (2005): Early depressive symptoms after stroke: Neuropsychological correlates and lesion characteristics. J Neurol Sci 228: 27–33.

47. Bruder GE, Stewart JW, McGrath PJ (2017, July 1): Right brain, left brain in depressive disorders: Clinical and theoretical implications of behavioral, electrophysiological and neuroimaging findings. Neuroscience and Biobehavioral Reviews, vol. 78. Elsevier Ltd, pp 178–191.

48. Weaver NA, Kuijf HJ, Aben HP, Abrigo J, Bae H-J, Barbay M, et al. (2021): Strategic infarct locations for post-stroke cognitive impairment: a pooled analysis of individual patient data from 12 acute ischaemic stroke cohorts. Lancet Neurol. https://doi.org/10.1016/S1474-4422(21)00060-0

