## Supplementary Material for "Strategic infarct locations for post-stroke depressive symptoms: a lesion- and disconnection-symptom mapping study"

***Magnetic Resonance Imaging***

Brain imaging was performed with a 3.0 tesla MRI scanner (Achieva, Philips Healthcare, Eindhoven, the Netherlands). Patients were scanned at hospital admission and at one-week follow-up; the follow-up scan was used for the current study. The MRI protocol included axial DWI and FLAIR sequences. DWI was obtained using an EPI-spin echo sequence with the following acquisition parameters: repetition time, 5,000 ms; echo time, 50 ms; diffusion b-value 1,000 s/mm2; slice thickness, 5 mm; intersection gap, 1 mm; matrix, 256 x 256; flip angle 90 degree. FLAIR imaging was obtained using a fast-spin echo sequence with the following acquisition parameters: repetition time, 11,000 ms; echo time, 125 ms; inversion time, 2,800 ms; slice thickness, 5 mm; intersection gap, 1 mm; matrix, 512 x 512; flip angle 90 degree.

***Definitions for impairment of ADL and cognitive impairment***

Impairment of Activities of Daily Living (ADL) was determined using the Korean Instrumental Activities of Daily Living (K-IADL) questionnaire (1). The K-IADL is comprised of 11 items: shopping, mode of transportation, ability to handle finances, housekeeping, preparing food, ability to use a telephone, responsibility for own medication, recent memory, hobbies, watching television, and fixing things around the house. Caregivers rated each item using a 4-point scale: 0=normal, 1=needs some assistance, 2=needs a lot of assistance, 3=impossible. Activities that were not performed before the onset of dementia were rated as “not applicable”. To obtain the total score, sum of scores was divided by the number of rated items except for “not applicable” items. ADL impairment was defined as a mean score of >0.43 (range: 0-3), based on validated cut-offs from a representative Korean population (1).

Post-stroke cognitive impairment (PSCI) was defined as cognitive impairment in one or more cognitive domains, in accordance with the VASCOG criteria for Vascular Cognitive Disorders (2). Cognitive performance was evaluated in the hospital setting using 60-minute Vascular Cognitive Impairment Harmonization Standards-Neuropsychology Protocol (3). Cognitive tests were assigned to five cognitive domains: attention and executive functioning; information processing speed; language; verbal memory; and visuospatial perception/construction. An overview of the categorization of cognitive tests is shown in Supplementary Table 6; further details were previously published (4). For each test performance, <5th percentile was defined as impaired. Performance on a cognitive domain was impaired if >50% of available neuropsychological tests on that domain were impaired, which was determined on a per-subject basis (i.e., patients might have a different number of available tests available per domain). Data on a minimum of three cognitive domains was needed to rule out PSCI.

***Validation sample selection and characteristics***

For the validation sample, patients were selected from the Hallym Vascular Cognitive Impairment (VCI) cohort, consisting of patients admitted to Hallym University Sacred Heart Hospital, Republic of Korea, with acute ischemic stroke between 2007 and 2018 (1). A total of 459 patients were selected based on the same inclusion criteria as the main sample from the Bundang VCI cohort. The only difference was that the 15-item Geriatric Depression Scale (GDS-15) assessment was used instead of the GDS-30. A flowchart of patient selection is provided in Supplementary Figure 1, and patient characteristics are shown in Supplementary Table 5.

The MRI protocol of the Hallym VCI cohort was similar to the protocol for the Bundang VCI cohort. The main difference was the timing of MRI assessment, which was performed after a median of 1 day for Hallym VCI (IQR: 1-2) and a median of 5 days for Bundang VCI (IQR: 4-6). The MRI protocols for both cohorts were previously described (4).

**Supplementary Table 1. Reviews on infarct location and post-stroke depression (2000-present)**

| Year | Reference | Methods | Inclusion period | N studies included | Conclusion – association with hemispheric lateralization of infarct? | Conclusion – association with infarct location? |
|---|---|---|---|---|---|---|
| 2017 | Zhang (5) | Meta-analysis | 1977 - 2016 | 31 | Yes, left hemisphere (only subacute phase) | N/A |
| 2017 | Douven (6) | Meta-analysis | 1989 - 2016 | 126 in qualitative analysis<br>74 in quantitative analysis | Acute infarcts: no association<br><br>Chronic infarcts: no association | Yes, frontal / basal ganglia |
| 2017 | Nickel (7) | Narrative review | N/A | N/A | No association | Yes, frontal pole / basal ganglia |
| 2016 | Robinson & Jorge (8) | Narrative review | N/A | N/A | N/A | Yes, left frontal / basal ganglia |
| 2015 | Wei (9) | Meta-analysis | 1977- 2013 | 43 | Acute infarcts: no association<br>Subacute infarcts: yes, right hemisphere | N/A |
| 2014 | Kutlubaev (10) | Meta-analysis of observational studies | 2004- 2013 | 23 | No association | N/A |
| 2006 | Spalletta (11) | Narrative review | N/A | 109 | Acute infarcts: yes, left hemisphere<br>Chronic infarcts: no associaton | Yes, prefrontal subcortical circuits |
| 2004 | Yu (12) | Meta-analysis | 1966- 2003 | 52 | Yes, right hemisphere | N/A |
| 2003 | Narushima (13) | Meta-analysis | 1981- 2000 | 27 | Acute infarcts: yes, left hemisphere | Yes, proximity to the frontal pole |
| 2000 | Carson (14) | Meta-analysis. Pooled relative risk | 1960- 1999 | 48 | No association | N/A |

**Supplementary Table 2. SVR-VLSM parameters**

| Scores | Optimized parameter C | Optimized epsilon | Prediction accuracy | P-value |
|---|---|---|---|---|
| Lesion-symptom mapping, corrected for age, sex, education | 0.001953125 | 0.05 | 0.40 | <0.0001 |
| Lesion-symptom mapping, corrected for age, sex, education, total infarct volume, NIHSS, ADL impairment and PSCI | 0.03125 | 0.01 | 0.57 | <0.0001 |
| Disconnection-symptom mapping, corrected for age, sex, education, total infarct volume, NIHSS, ADL impairment and PSCI | 0.03125 | 0.05 | 0.48 | <0.0001 |

**Supplementary Table 3. SVR-ROI parameters**

| Scores | Optimized parameter C | Prediction accuracy | P-value |
|---|---|---|---|
| Lesion-symptom mapping, corrected for age, sex, education | 0.00391 | 0.15 | 0.21 |
| Lesion-symptom mapping, corrected for age, sex, education, total infarct volume, NIHSS, ADL impairment and PSCI | 0.00391 | 0.11 | 0.29 |

**Supplementary Table 4. Significant voxels from SVR-VLSM results per region of interest**

| ROI | N voxels in ROI | N significant voxels from SVR-LSM | % of ROI |
|---|---|---|---|
| Right Amygdala | 1863 | 157 | 8.4 |
| Right Pallidum | 2209 | 163 | 7.4 |
| Right Hippocampus | 7280 | 374 | 5.1 |
| Right Posterior corona radiata | 5953 | 271 | 4.6 |
| Right Sagittal stratum | 2173 | 81 | 3.7 |
| Left Cingulum of the hippocampus | 1650 | 53 | 3.2 |
| Right Fornix (cres) / Stria terminalis | 1105 | 31 | 2.8 |
| Left Fornix (cres) / Stria terminalis | 1131 | 31 | 2.7 |
| Right Tapetum | 663 | 18 | 2.7 |
| Left Hippocampus | 7239 | 170 | 2.3 |
| Right Posterior thalamic radiation | 5400 | 101 | 1.9 |
| Right Posterior limb of the internal capsule | 2492 | 45 | 1.8 |
| Right Retrolenticular part of the internal capsule | 2666 | 42 | 1.6 |
| Right Temporal inferior gyrus | 28002 | 314 | 1.1 |
| Left Cingulum (cingulate.gyrus) | 3670 | 41 | 1.1 |
| Right Superior longitudinal fasciculus | 9580 | 102 | 1.1 |
| Left ParaHippocampal gyrus | 7644 | 81 | 1.1 |

The number of significant voxels from the primary SVR-VLSM analysis ($p<0.05$; shown in red in main Figure 1C) was calculated for each region of interest (ROI) from the AAL and ICBM-DTI-81 atlases. Only ROIs with at least 1% significant voxels are shown. ROIs are listed in descending order of percentage of significant voxels within the ROI.

**Supplementary Table 5. Clinical characteristics of main sample and validation sample**

| **Demographics and clinical characteristics** | **Bundang VCI (N=553)** | **Hallym VCI (N=459)** |
|---|---|---|
| Age (years), mean (SD) | 69.0 (11.0) | 64.9 (12.0) |
| Female, N (%) | 233 (42%) | 196 (43%) |
| Years of education, median (IQR) | 9 (6-14) | 9 (6-12) |
| NIHSS at admission, median (IQR) | 3 (1-5) | 2 (1-4) (missing N=24) |
| K-IADL score, median (IQR) | 0.10 (0.00-0.42), (missing N=4) | 0.00 (0.00-0.20) (missing N=24) |
| ADL impairment (K-IADL score > 0.43) | 137 (25%) (missing N=4) | 61 (13%) (missing N=24) |
| **Hand preference, N (%)** | (missing N=2) | (missing N=17) |
| Right | 534 (97%) | 435 (98%) |
| Left | 6 (1%) | 4 (1%) |
| Ambidextrous | 11 (2%) | 3 (1%) |
| **Vascular risk factors, N (%)** | | |
| Hypertension | 422 (76%) | 259 (57%) (missing N=1) |
| Hyperlipidemia | 135 (24%) | 166 (37%) (missing N=8) |
| Current smoker | 119 (22%) | 126 (28%) (missing N=8) |
| Past smoker | 113 (20%) | 43 (9%), (missing N=17) |
| Diabetes Mellitus | 179 (32%) | 139 (30%) (missing N=1) |
| Atrial fibrillation | 84 (15%) (missing N=27) | 44 (10%) (missing N=7) |
| **Neuropsychiatric and cognitive assessment** | | |
| Time interval between stroke onset and neuropsychiatric and cognitive assessment (days), median (IQR), range | 105 (11-168), range 1-361 | 97 (90-104), range 5-307 |
| GDS-30 scores, median (IQR), range | 13 (7-20), range 0-30 | N/A |
| GDS-15 scores, median (IQR), range | N/A | 4 (1-8), range 0-15 |

| | | |
|---|---|---|
| Presence of post-stroke cognitive impairment, N (%) | 323 (58%)<br>(missing N=10) | 200 (44%) |
| **Brain MRI** | | |
| Time interval between stroke onset and MRI (days), median (IQR), range | 5 (4-6), range 0-41 | 1 (1-2), range 0-15 |
| Total infarct volume in ml[‖], median (IQR), range | 3.3 (1.1-15.1),<br>range 0.04-535.1 | 1.9 (0.9-9.0),<br>range 0.06-247.5) |

If any data was missing, this is noted behind the respective variable. Valid percent is indicated in cases with missing data. Abbreviations: IQR, interquartile range; MRI, magnetic resonance imaging; SD, standard deviation.

**Supplementary Table 6. Categorization of cognitive tests across cognitive domains for determining post-stroke cognitive impairment**

| | **Attention and executive functions** |
|---|---|
| **Attention and executive functions** | Phonemic fluency (three phonemes, number of words in one minute per phoneme) (15) |
| | Korean-Trail Making Test - Elderly's version B (time in seconds) (16) |
| **Language** | Short Form of the Korean-Boston Naming Test (number correct) (17) |
| | Semantic fluency, category animals (number of words in one minute) (15) |
| **Processing speed** | Korean-Trail Making Test - Elderly's version A (time in seconds) (16) |
| | Digit Symbol Coding (number correct) (18) |
| **Verbal memory**† | Seoul Verbal Learning Test - immediate recall (number correct) (19) |
| | Seoul Verbal Learning Test – delayed recall (number correct) (19) |
| | Seoul Verbal Learning Test – recognition (number correct) (19) |
| **Visuoperception and -construction** | Rey Complex Figure Test: Copy task (19) |

† Each of the Seoul Verbal Learning Test scores was included as a separate test for the verbal memory domain. Abbreviation: SD, standard deviation.

**Supplementary Figure 1. Flow chart of patient selection for validation sample**

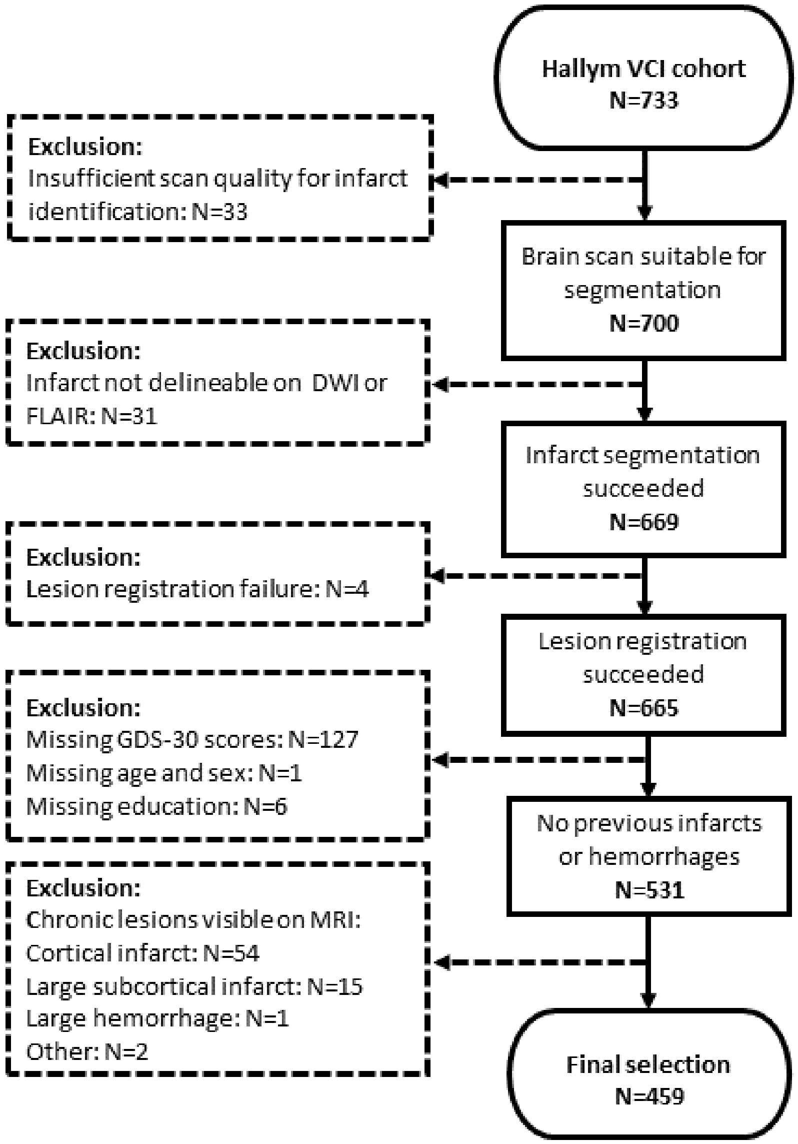

**Supplementary Figure 2. Comparison of lesion prevalence in main sample and validation sample**

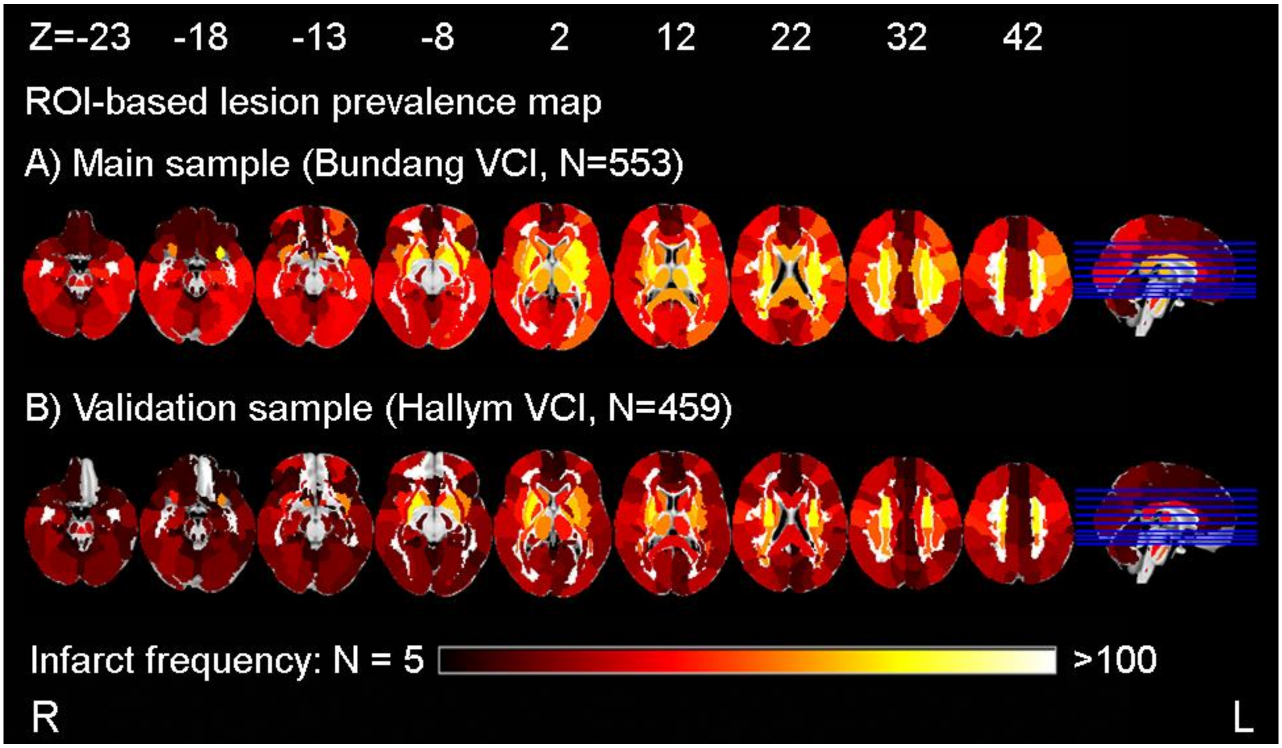


This figure shows a direct comparison of the lesion distribution and brain lesion coverage in the main sample (A) and validation sample (B). In order to perform external validation of selected regions of interest (ROIs), a sufficient number of patients need to have infarcts in these ROIs to achieve sufficient statistical power. A minimum of 5 patients was required in the current study. Colors indicate the lesion prevalence at the level of regions of interest (ROI). Overall lesion prevalence is lower, yet nearly all ROIs were still damaged in 5 or more patients and therefore eligible for analysis. Of note, for the three selected ROIs (i.e. right amygdala, right hippocampus and right pallidum), and the three negative control ROIs, coverage was comparable in both datasets (see Table 2 in main text).

**Supplementary References**


1. Kang SJ, Hye Choi S, Lee BH, Kwon JC, Na DL, Han S-H (2002): The Reliability and Validity of the Korean Instrumental Activities of Daily Living (K-IADL). *J Korean Neurol Assoc*, vol. 20.
2. Sachdev P, Kalaria R, O'Brien J, Skoog I, Alladi S, Black SE, *et al.* (2014): Diagnostic criteria for vascular cognitive disorders: A VASCOG statement. *Alzheimer Dis Assoc Disord* 28: 206–218.
3. Yu KH, Cho SJ, Oh MS, Jung S, Lee JH, Shin JH, *et al.* (2013): Cognitive impairment evaluated with vascular cognitive impairment harmonization standards in a multicenter prospective stroke cohort in Korea. *Stroke* 44: 786–788.
4. Weaver NA, Kancheva AK, Lim J-S, Biesbroek JM, Wajer IMH, Kang Y, *et al.* (2021): Post-stroke cognitive impairment on the Mini-Mental State Examination primarily relates to left middle cerebral artery infarcts. *Int J Stroke* 174749302098455.
5. Zhang Y, Zhao H, Fang Y, Wang S, Zhou H (2017): The association between lesion location, sex and poststroke depression: Meta-analysis. *Brain Behav* 7: 1–11.
6. Douven E, Köhler S, Rodriguez MMF, Staals J, Verhey FRJ, Aalten P (2017, September 1): Imaging Markers of Post-Stroke Depression and Apathy: a Systematic Review and Meta-Analysis. *Neuropsychology Review*, vol. 27. Springer New York LLC, pp 202–219.
7. Nickel A, Thomalla G (2017, September 21): Post-stroke depression: Impact of lesion location and methodological limitations-a topical review. *Frontiers in Neurology*, vol. 8. Frontiers Media S.A. https://doi.org/10.3389/fneur.2017.00498
8. Robinson RG, Jorge RE (2016): Post-stroke depression: A review. *Am J Psychiatry* 173: 221–231.
9. Wei N, Yong W, Li X, Zhou Y, Deng M, Zhu H, Jin H (2015): Post-stroke depression and lesion location: A systematic review. *J Neurol* 262: 81–90.
10. Kutlubaev MA, Hackett ML (2014): Part II: Predictors of depression after stroke and impact of depression on stroke outcome: An updated systematic review of observational studies. *Int J Stroke* 9: 1026–1036.

11. Spalletta G, Bossù P, Ciaramella A, Bria P, Caltagirone C, Robinson RG (2006): The etiology of poststroke depression: A review of the literature and a new hypothesis involving inflammatory cytokines. *Mol Psychiatry* 11: 984–991.

12. Yu L, Liu CK, Chen JW, Wang SY, Wu YH, Yu SH (2004): Relationship between post-stroke depression and lesion location: A meta-analysis. *Kaohsiung Journal of Medical Sciences*, vol. 20. Elsevier (Singapore) Pte Ltd, pp 372–380.

13. Narushima K, Kosier JT, Robinson RG (2003): A Reappraisal of Poststroke Depression, Intra- and Inter-Hemispheric Lesion Location Using Meta-Analysis. *J Neuropsychiatry Clin Neurosci* 15: 422–430.

14. Carson AJ, MacHale S, Allen K, Lawrie SM, Dennis M, House A, Sharpe M (2000): Depression after stroke and lesion location: A systematic review. *Lancet* 356: 122–126.

15. Kang, Y., Chin, J. H., Na, D. L., Lee, J., & Park JS (2000): A normative study of the Korean version of Controlled Oral Word Association Test (COWAT) in the elderly. *Korean J Clin Psychol*.

16. Yi H, Chin J, Lee B, Kang Y (2007): Development and validation of Korean version of Trail Making Test for elderly persons. *Dement Neurocognitive Disord*. 6.2: 54-66.

17. Kang Y, Kim H, Psychol DN-KJC (1999): A short form of the Korean-Boston Naming Test (K-BNT) for using in dementia patients. *Korean J Clin Psychol.* 18: 125-138.

18. Yum T, Park Y, Oh-hashi K, Kim J, Lee Y, Yum T (1992): The manual of Korean-Wechsler adult intelligence scale.

19. Kang Y, Na D, Hahn S (2003): Seoul neuropsychological screening battery. *Incheon: Human brain research & consulting co*.
